# Virtual drug screen reveals context-dependent inhibition of cardiomyocyte hypertrophy

**DOI:** 10.1101/2022.08.22.504776

**Authors:** Taylor G. Eggertsen, Jeffrey J. Saucerman

## Abstract

**Background and Purpose:** Pathological cardiomyocyte hypertrophy is a response to cardiac stress that typically leads to heart failure. Despite being a primary contributor to pathological cardiac remodeling, the therapeutic space that targets hypertrophy is limited. Here, we apply a network model to virtually screen for FDA-approved drugs that induce or suppress cardiomyocyte hypertrophy.

**Experimental Approach:** A logic-based differential equation model of cardiomyocyte signaling was used to predict drugs that modulate hypertrophy. These predictions were validated against curated experiments from the prior literature. The actions of midostaurin were validated in new experiments using TGFβ- and NE-induced hypertrophy in neonatal rat cardiomyocytes.

**Key Results:** Model predictions were validated in 60 out of 70 independent experiments from the literature and identify 38 inhibitors of hypertrophy. We additionally predict that the efficacy of drugs that inhibit cardiomyocyte hypertrophy is often context dependent. We predicted that midostaurin inhibits cardiomyocyte hypertrophy induced by TGFβ, but not NE, exhibiting context dependence. We further validated this prediction by *in vitro* experimentation. Network analysis predicted critical roles for the PI3K and RAS pathways in the activity of celecoxib and midostaurin, respectively. We further investigated the polypharmacology and combinatorial pharmacology of drugs. Brigatinib and irbesartan in combination were predicted to synergistically inhibit cardiomyocyte hypertrophy.

**Conclusion and Implications:** This study provides a well-validated platform for investigating the efficacy of drugs on cardiomyocyte hypertrophy, and identifies midostaurin for consideration as an antihypertrophic drug.

**‘What is already known’:**

- Cardiac hypertrophy is a leading predictor of heart failure.
- Cardiomyocyte hypertrophy is driven by intracellular signaling pathways that are not targeted by current drugs

**‘What this study adds’:**

- Computational model integrates 69 unique drugs to predict cardiomyocyte hypertrophy
- Drug-induced inhibition of cardiomyocyte hypertrophy is context-dependent
- Midostaurin inhibits TGFβ-induced cardiomyocyte hypertrophy

**‘Clinical significance’:**

- Midostaurin is identified as a candidate antihypertrophic drug
- Several FDA approved drugs are predicted to inhibit cardiomyocyte hypertrophy either individually or in combination.

## INTRODUCTION

Heart failure remains one of the most critical health and economic concerns^1^. Hypertrophy is a leading predictor of heart failure^2–6^, and a good therapeutic target for mitigating onset of heart failure^7,8^. Understanding the signaling pathways in hypertrophy is critical to identifying drug targets^9^. Current drugs still do not meet the growing demands of heart failure. Current therapeutics include ACE inhibitors, Beta blockers, ARBs, diuretics, hydralazine and nitrates^10^. These target only the alpha-adrenergic and beta-adrenergic receptors or mitigate hypertension via vasodilation or alleviating water retention. Recent studies have also identified SGLT2 inhibitors as promising therapeutics, however their mechanism in heart failure is not fully understood^11^. There are still many unexplored intracellular targets in hypertrophic signaling. Understanding the mechanisms of pathological hypertrophy paves the way for therapeutic development.

Cardiac hypertrophy signaling is very complex, with much crosstalk between pathways. The challenge of integrating current knowledge of hypertrophic signaling can be met with computational modeling approaches^12^. We previously developed and validated large network computational models that simulate cardiomyocyte hypertrophy signaling^13^, cardiac fibroblast signaling^14^, and mechanosignaling^15^. Therapeutics have not been modeled in the context of cardiomyocyte hypertrophy, however. Here we develop a drug-target model of cardiomyocyte signaling to predict the effect of FDA approved drugs on hypertrophy. We use sensitivity analysis to identify potential network mechanisms of drug activity. Using these predictions, we develop and experimentally validate model-guided hypotheses. Finally, we examined the effects of both polypharmacology and drug combinations on cardiomyocyte hypertrophy. These studies provide a foundation for systems pharmacology of cardiac remodeling.

## METHODS

### Signaling model

Cardiomyocyte signaling was modeled using a previously published logic-based differential equation (LDE) formalism^13^. We used the Netflux software (https://github.com/saucermanlab/Netflux) to construct LDE models from a literature-based network of nodes. This network model is composed of 107 nodes, with 17 biochemical inputs including 1 mechanical stretch input. Outputs include hypertrophic markers bMHC, ANP, BNP, and cell area. These LDEs were then solved in MATLAB using the solver ode15s. Values of 0.02 were set for all input weights and the system was run to steady state to establish baseline output values. Biochemical stimulation was simulated by setting one of the inputs to a value of 0.1, resulting in 17 possible hypertrophic environments.

### Drug simulations

Drug characteristics were retrieved from the DrugBank database, which maintains pharmacokinetic and drug target data on over 7000 FDA approved or clinically investigational drugs^16^. Drugs that target nodes within the network model were identified from this database. A program was developed to extract drug agonism data for each compound along with their name, database ID, and listed drug target. Drug binding properties were manually curated from PubMed. Through this method 258 drugs were identified that target nodes in the network model, many of which share drug targets. From this group, 69 unique drug-target interactions were further identified. Each unique drug-target interaction consists of a unique drug, binding, agonism and target combination.

Drug activity was implemented into the model using a revision of the ODE file produced from Netflux. Drugs were split into groups defined by binding properties (competitive or non-competitive) and action (agonist or antagonist). The binding properties determine how the upstream node activity is shifted by the drug dose, while the action determines whether the node is upregulated (agonist) or downregulated (antagonist). These changes impact the weight parameter of the affected node, which is then saved as an updated parameter. The equations that govern these dynamics were developed previously by Zeigler et al^17^. Each simulation is run first without drug to establish a baseline activity, and then with the appropriate drug to identify changes in activity. All drugs were simulated at 80% of saturated dose (**Figure S1**).

Drug pair simulations were performed by implementing two drugs into the model and combining the changed parameters. Synergy scores were calculated by subtracting the Bliss predicted inhibition rate y_ab_ from the model’s predicted inhibition. The value y_ab_ is calculated using the equation:

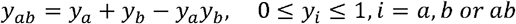

where y_a_ and y_b_ are the inhibition rates with drug A alone at dose a or drug B alone at dose b.

### Literature validation

To validate the predictions of drug simulation we performed manual literature curation of in vitro and in vivo rat and mouse experiments not used to build the model. We collected results from 38 experiments in which FDA approved drugs were used to inhibit cardiac hypertrophy both in vitro and in vivo. We additionally identified a screen of 3241 drugs to identify those which inhibit phenylephrine induced cardiomyocyte hypertrophy in vitro^18^. The conditions in these experiments were simulated in the model by implementing the drug used, or a drug in the same class, in the same hypertrophic environment used to stimulate the cells or animal. The threshold for validating a given prediction is based on a 0.1% change in activity which was used in previous publications^13,19^. In total 59 *in vitro* findings and 11 *in vivo* experiments were used to validate the model.

### Mechanistic subnetworks

We performed sensitivity analysis to identify the nodes in the network that mediate the effect of each drug. Knocking down individual nodes in the presence of stimulus describes the role of the nodes in the network for that stimulus. Knocking down individual nodes in the presence of stimulus plus drug describes the role of the nodes during drug activity. The difference between these node responses identifies the nodes that are critical for mediating drug activity. These critical nodes identified by sensitivity analysis are then compared to nodes that are affected by the drug. The intersection of these node groups results in a subnetwork that can be used as a mechanistic map of the drug. These subnetworks were visualized using Cytoscape^20^.

### In vitro validation

Neonatal cardiomocytes were isolated from 1-2-day old Sprague-Dawley rats using the Neomyts isolation kit (Cellutron, Baltimore MD). Cardiomyocytes were cultured in plating media (Dulbecco Modified Eagle Media, 17% M199, 10% Horse Serum, 5% Fetal bovine Serum, 100 U/ml penicillin, and 50 mg/ml streptomycin) in 96 well plates, pretreated with SureCoat, at a density of 30,000 cells/well. 48 hours post isolation, cardiomyocytes were changed to serum-free maintenance media (Dulbecco Modified Eagle Media, 19% M199, 1% ITSS, 100 U/ml penicillin, and 50 mg/ml streptomycin) for 24 hours. Cardiomyocytes were then treated with one of three hypertrophic stimuli (200 nM angiotensin II, 5 μM norepinephrine (NE), 5 ng/ml transforming growth factor β (TGFβ)), 10% FBS, or negative control. The cells were simultaneously treated with specified concentrations of midostaurin. The cardiomyocytes were left treated for 48 hours, at which point they were fixed with 4% paraformaldehyde for 20 minutes.

Cardiomyocytes were permeabilized with 0.1% Triton-X for 15 minutes. Cardiomyocytes were blocked with 1% bovine serum albumin in PBS for 1 hour, then treated with mouse anti-α-actinin primary antibody at a concentration of 1:200 overnight. Cardiomyocytes were then blocked with a 5% goat serum in PBS for 1 hour, then Alexa Fluor-568-conjugated goat anti-mouse secondary antibody at a concentration of 1:200 was applied for 1 hour. The cells were stained with DAPI prior to imaging.

High-content imaging was performed on the stained cardiomyocytes using an Operetta CLS High Content Analysis System courtesy of Mohammad Fallahi-Sichani. These images were processed using CellProfiler^21^, in which an algorithm developed previously^22^ was used to identify nuclei and cell borders. Cell area was measured for each cell in all conditions, and cells with undetectable cytoplasms were not counted.

### Statistics

Statistical significance between conditions, considering two separate cellular isolations, was determined by a 2-way ANOVA followed by Dunnet’s test for multiple comparisons, where each drug condition was compared to the hypertrophic control without drug.

## RESULTS

### Virtual screen identifies drugs that stimulate and inhibit cardiomyocyte hypertrophy

To simulate the effects of drugs on cardiac hypertrophy, we implemented drug activity into a logic-based signaling network. This network model was previously manually curated to represent signaling molecules as nodes and directed interactions as edges^13,23^. Here, drugs that target the nodes of this network were identified from the DrugBank database (**Figure 1A**). Out of over 7000 FDA-approved or investigational pharmaceuticals, this pipeline identified 258 drugs that directly target proteins in the network, with 69 unique drug-target pairings.

**Figure 1.**
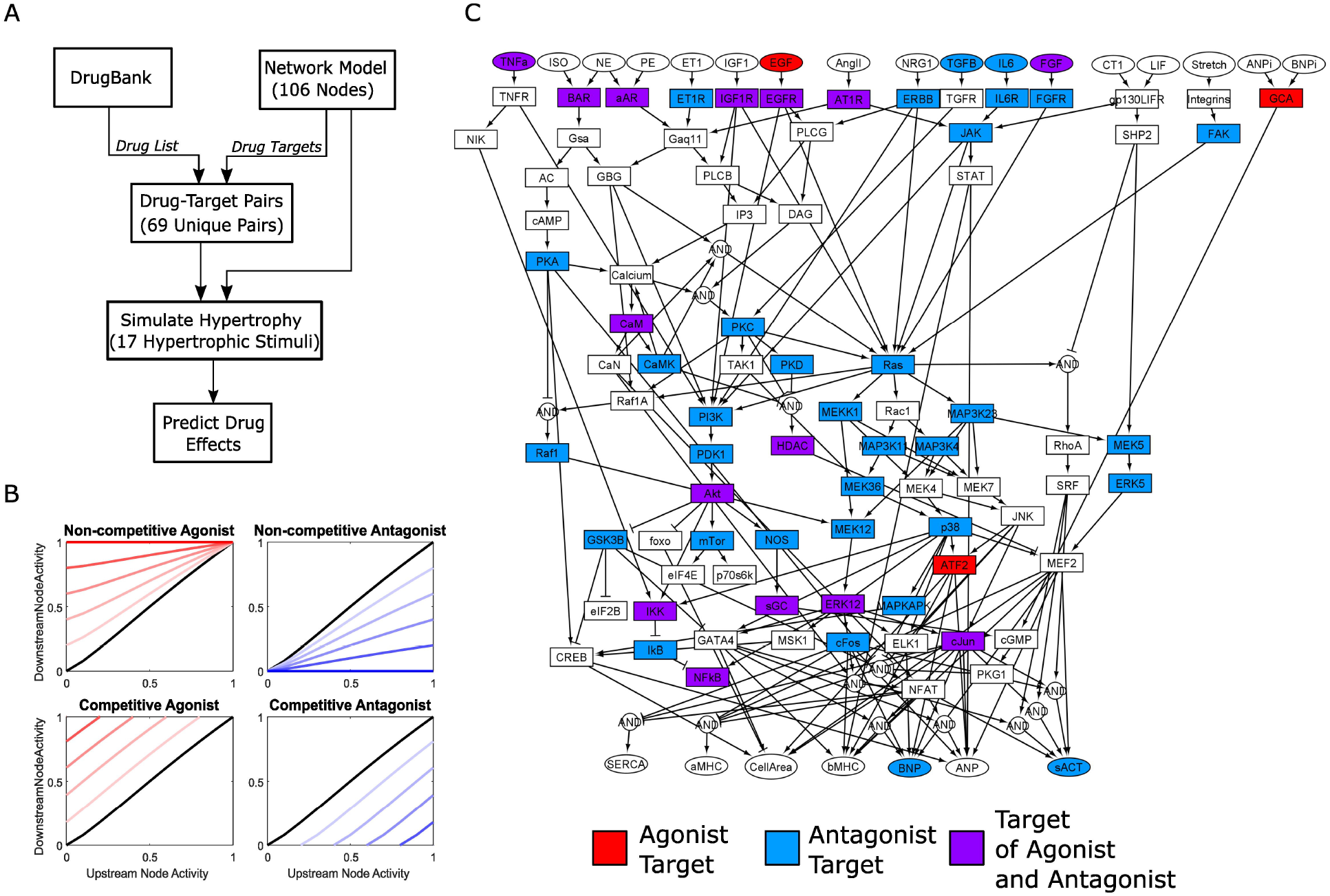
A curated signaling network model identifies drugs that may be repurposed to treat cardiac hypertrophy. A) Schematic of an in silico drug testing pipeline showing the implementation of drugs from the DrugBank database into the hypertrophy signaling network. B) Drugs are classified primarily as either agonists (red) or antagonists (blue). Drugs are further divided into non-competitive and competitive categories, which dictate how downstream node activity is determined from the upstream signal. Increasing drug concentration is indicated by darker colored lines, while the no drug control is indicated by the black lines. C) Drug targets in the hypertrophy signaling network exhibit high coverage of the network space, with 69 unique drug-target pairs.

The effect of a drug on the activity downstream of the targeted node is dependent on whether the drug is agonistic or antagonistic and binds competitively or non-competitively (**Figure 1B**). In general, the agonism of the drug determines whether the signal is decreased or increased compared to normal, and the binding of the drug determines the slope of the node activity. The targeted nodes span across the entire network (**Figure 1C**). Most drugs target a single node, but there are 24 representative drugs that target multiple nodes (**Table 1, Table S1**). In addition, several of the nodes are targeted by multiple drugs. Unique drug-target pairs are representative of similar drugs acting on the same target(s). Much of the network is targeted by drugs not traditionally used in cardiovascular treatment.

**Table 1.**
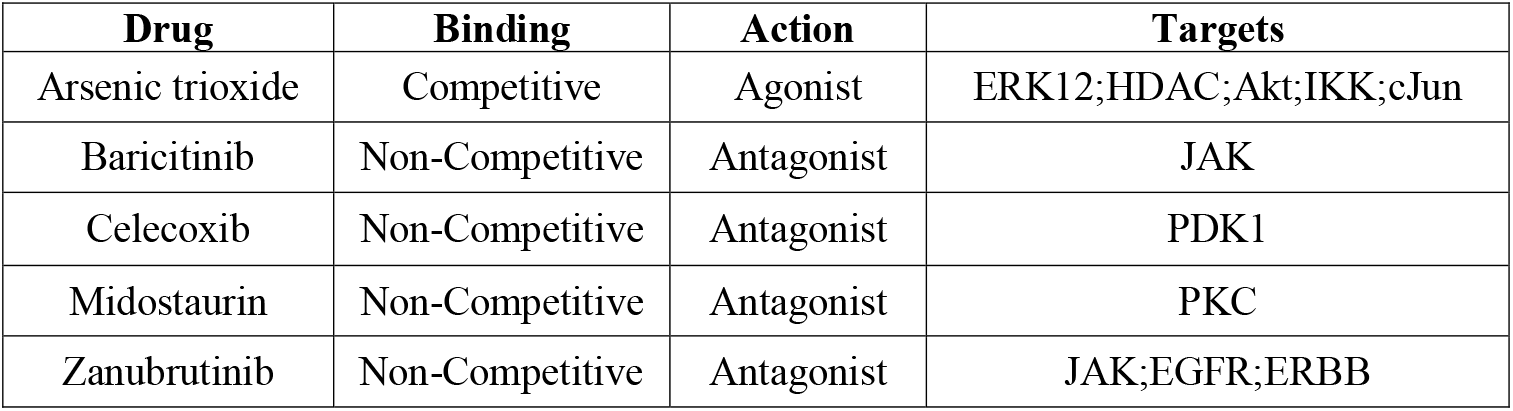
Representative drugs identified from the virtual screen along with their characteristics

A virtual screen was performed on the identified drugs to determine their effect on hypertrophy. Drugs that share agonism, binding, and targets are represented by a single drug in the output of the virtual screen. These drugs are shown to increase or decrease the activity of their appropriate targets (**Figure S1**). In total 38 out of 69 unique drug-target pairs were identified from the virtual screen that inhibit hypertrophy, with an additional 14 unique drug-target pairs that stimulate hypertrophy (Table S1).

The virtual screen was then used to identify the effect of different hypertrophic environments on the drug inhibition or stimulation of hypertrophy. The network was first stimulated by one of 17 hypertrophic stimuli, and then the activity of the drug was simulated. The predicted behavior of the cell in response to these drugs was determined from the 7 phenotypic outputs of the model, which include cell area and fetal gene expression. Phenotypic outputs exhibit fairly consistent patterns for the representative drugs (**Figure S2**), so we focused subsequent analysis on cell area.

### Drugs modulate hypertrophy in a context-dependent manner

Simulations of drug activity reveal varying drug efficacy across 17 hypertrophic stimulid. Some drugs exhibited context-independent induction of hypertrophy (e.g. arsenic trioxide, atorvastatin) or inhibition of hypertrophy (e.g. baricitinib, zanubrutinib) despite variation in hypertrophic stimulus (**Figure 2A**). This may indicate that these drugs impact critical network hubs that integrate multiple signaling pathways. In contrast, other drugs exhibited context-specific effects, inhibiting only with selected hypertrophic stimuli. For example, celecoxib is effective in NE-induced hypertrophy while midostaurin is effective only in TGFβ-induced hypertrophy (**Figure 2A**). This suggests that these drugs stimulate hypertrophic signaling through different pathways in the network model.

**Figure 2.**
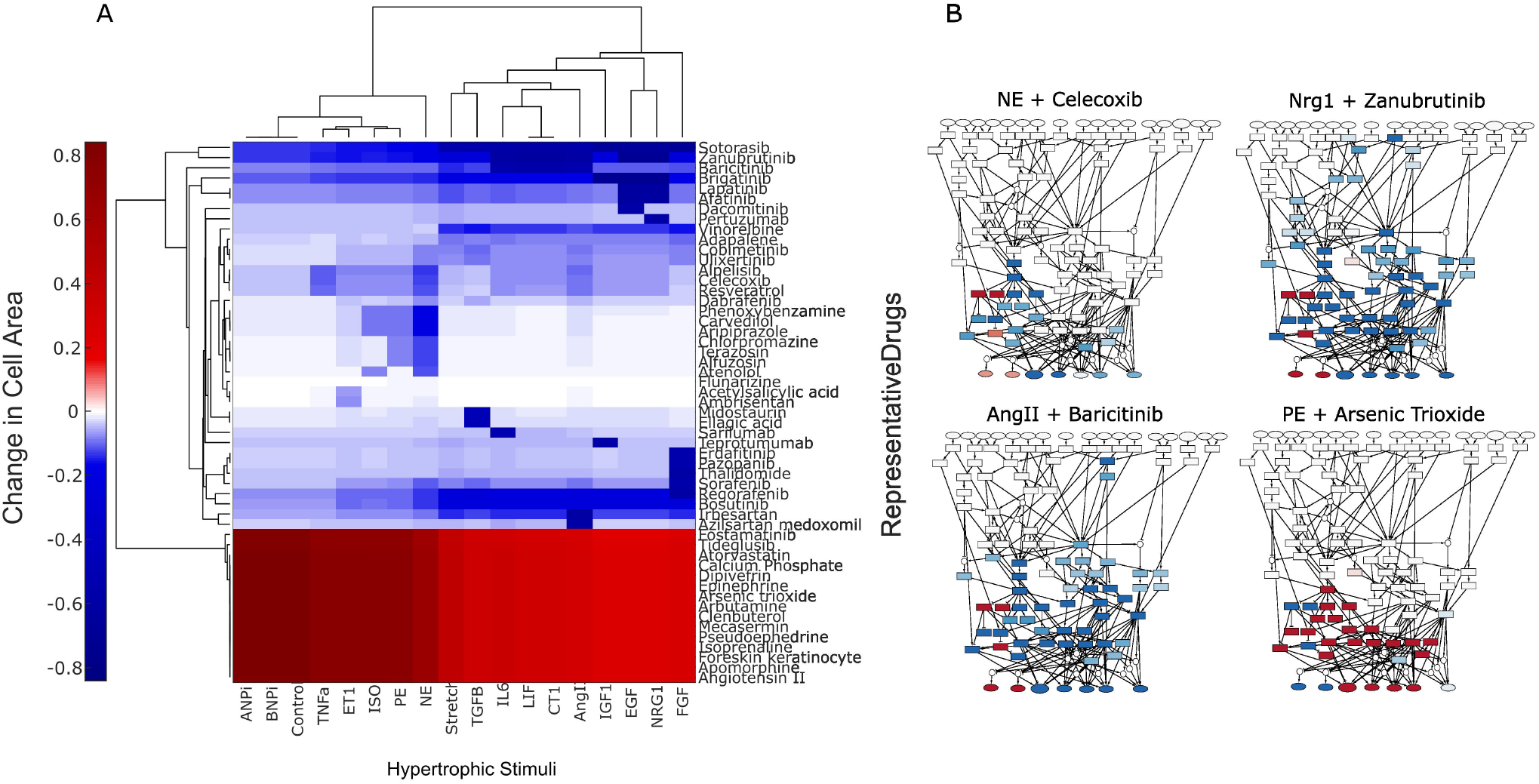
Simulation of drug activity predicts context-dependent modulation of hypertrophy. A) The predicted change in cell area is shown for each representative drug across simulations of 17 biochemical environments. Each row shows the predicted effect for one of 52 unique drugs on cell area. Each column indicates the role that the hypertrophic stimulus plays in the activity of the 52 drugs. B) Selected whole-network maps showing the change in activity of network nodes in response to the given combination of drug and hypertrophic stimulus.

As visualized in **Figure 2B**, the network response to celecoxib, baricitinib, zanubrutinib, and arsenic trioxide is highly dependent on both the hypertrophic stimulus and the particular drug implemented. Drugs that target nodes further downstream in the network, like celecoxib or arsenic trioxide, reveal a much more localized response from the network. Drugs that target nodes further upstream, like baricitinib or zanubrutinib, show activity change across much more of the network. Drugs that inhibit hypertrophy, such as celecoxib, zanubrutinib, and baricitinib, can be seen to decrease node activity along critical pathways in the network. Drugs that stimulate hypertrophy, such as arsenic trioxide, are shown to increase node activity in the network.

We next evaluated whether the model predictions were consistent with experimental data from prior literature, which are independent from the data used to develop the model. The model is successfully validated in 32 out of 38 (84%) individual hypertrophy experiments, all performed in cardiomyocytes from mice or rats^24–53^ (**Figure 3A**). The model additionally is validated in 28 out of 32 (88%) outcomes from an in vitro drug screen of phenylephrine-induced cardiomyocyte hypertrophy^18^ (**Figure 3B**). Out of these 70 independent experiments, 11 were performed *in vivo* with 91% agreement, and 59 were performed *in vitro* with 85% agreement. The model retained robustness in experimental validation when varying the validation threshold up to 5% (**Figure S3**). Overall, the pharmacological model of cardiomyocyte hypertrophy is 86% predictive of independent experimental results.

**Figure 3.**
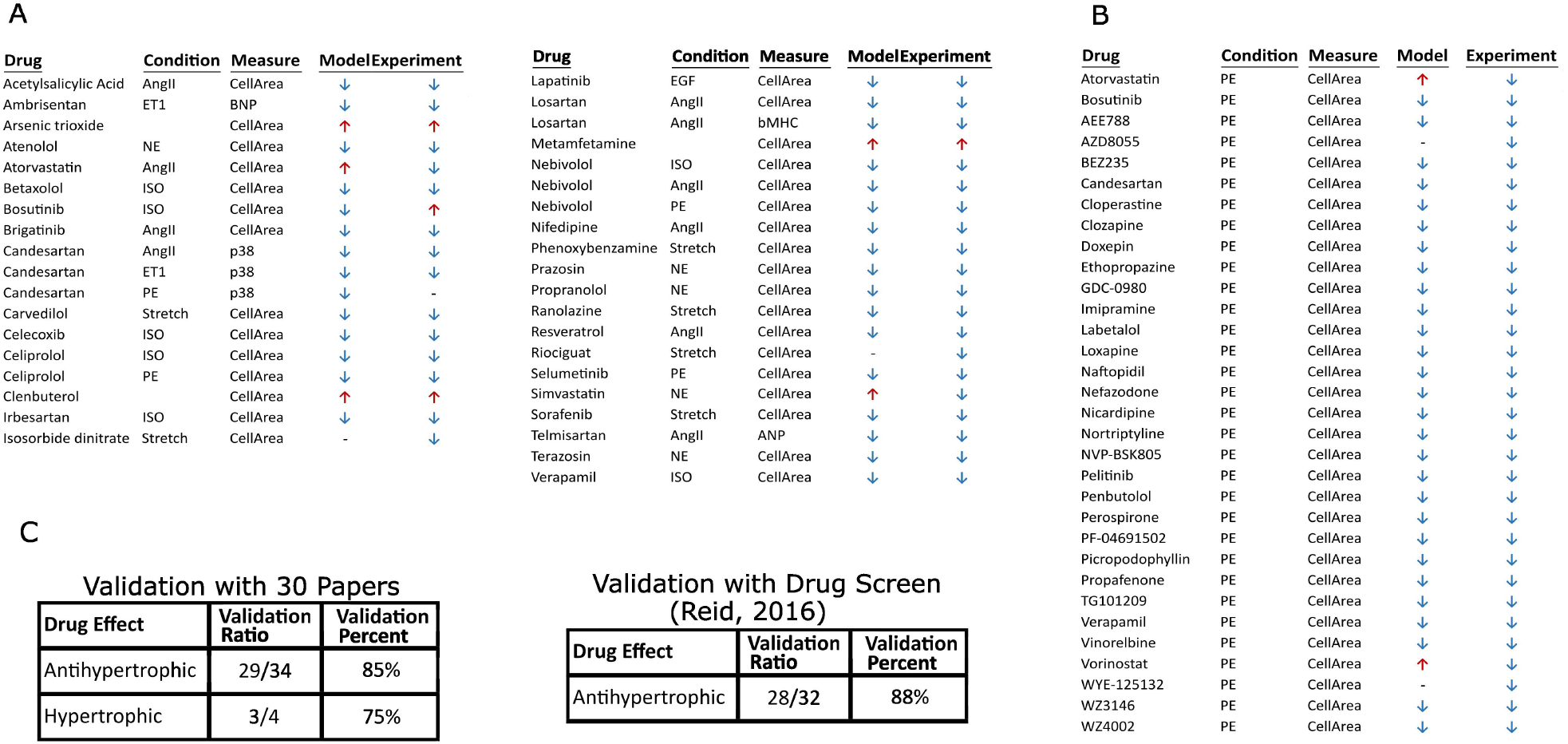
Predicted effects of drug activity agree well with prior experimental literature. A) Model validation predicts the phenotype of 32 of 38 experiment results from both in vitro and in vivo cardiomyocyte hypertrophy studies from the literature. B) Model validation predicts 28 of 32 experimental results from an in vitro drug screen of phenylephrine induced hypertrophy (PMID: 27130278). C) Summary of validation results show over 80% validation when predicting antihypertrophic drug effects.

### Mechanisms of action are predicted for anti-hypertrophic drugs

Next, we performed sensitivity analysis to investigate the network mechanism of action of the drugs. Each node of the network was individually knocked down to determine its impact on cell area in the presence of a hypertrophic stimulus (**Figure S4**). This analysis was performed with and without the presence of drug, and the difference between these was used to identify the nodes that mediate drug activity. These mediator nodes were intersected with those whose signaling activity was impacted by the drug, resulting in a subnetwork that represents a predicted network mechanism of action for a given drug (**Figure 4**).

**Figure 4.**
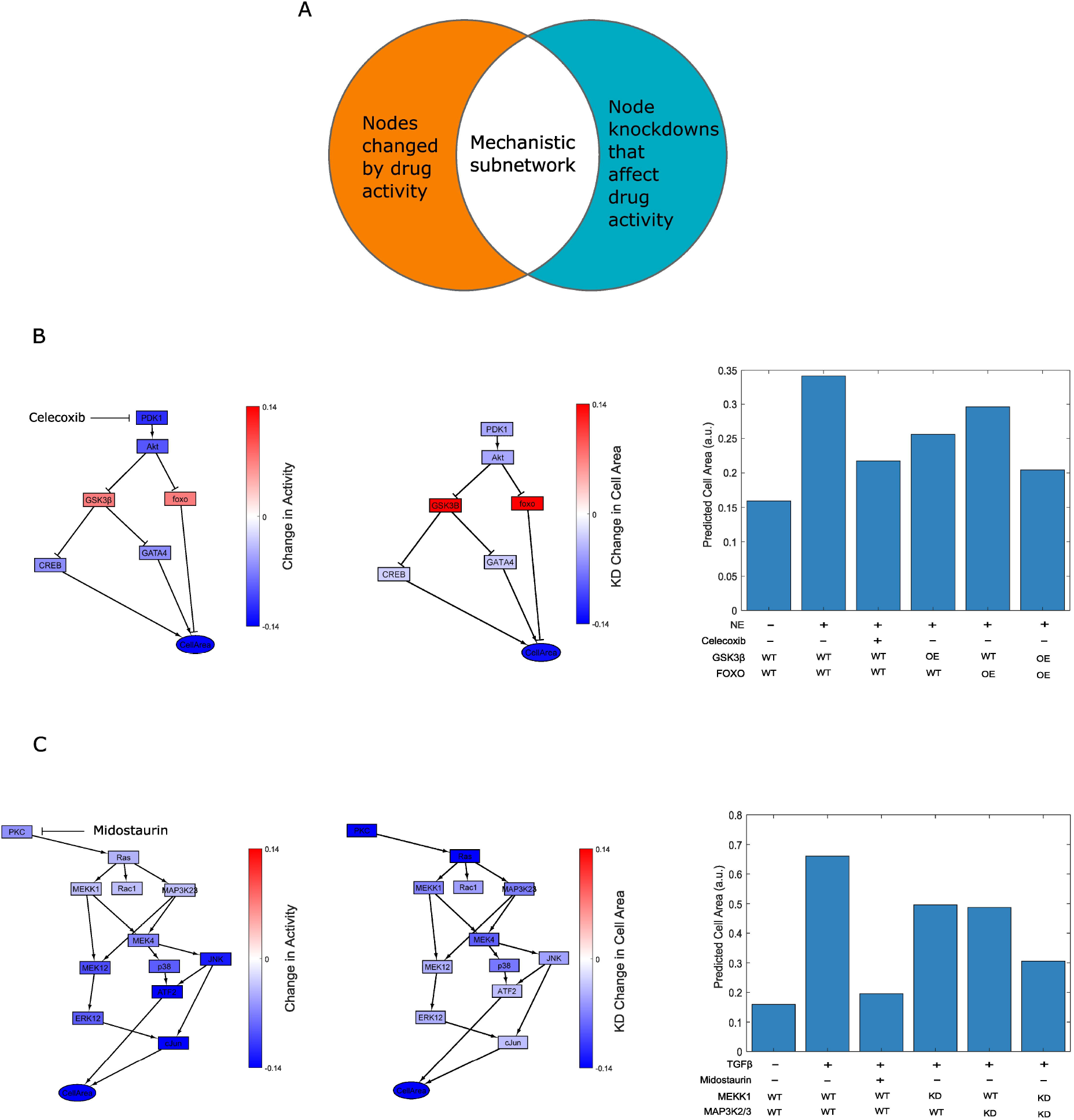
Subnetworks predict mechanisms by which drugs inhibit hypertrophy in the contexts of NE, or AngII stimulation. A) The mechanistic subnetwork is the intersection of nodes that are altered by drug activity, together with nodes whose knockdown modulate drug activity. B) Celecoxib inhibits NE-induced hypertrophy by derepressing both GSK3β and foxo. C) Midostaurin inhibits TGFβ-induced hypertrophy by repressing RAS through MAP3K1 and MAP3K2/3. WT, wild type; OE, overexpression; KD, knockdown

Celecoxib is a cyclo-oxygenase (COX-2) inhibitor as well as a weak inhibitor of 3-phosphoinositide-dependent kinase-1 (PDK1). Despite evidence of hypertrophic inhibition, the mechanism of action of celecoxib in hypertrophy is still uncertain^32,54,55^. Mechanistic subnetwork analysis predicted that celecoxib inhibits norepinephrine-induced hypertrophy via PDK1 and parallel GSK3B/FOXO pathways. Overexpression of these two nodes results in inhibition of NE-induced hypertrophy in a manner comparable to the action of celecoxib (**Figure 4B**). Although celecoxib is the only PDK1 inhibitor identified in the screen, these predictions would extend to other PDK1 inhibitors with similar binding characteristics.

Midostaurin is a FLT3 inhibitor which has been shown to inhibit several receptor tyrosine kinases, including KIT, PDGFRα/β and members of the PKC family. Although some evidence suggests FLT3 inhibitors show cardiotoxic effects, safety evaluations demonstrate that midostaurin does not cause QT prolongation as seen in other FLT3 inhibitors^56^. Hypertension and pericardial effusion are the primary cardiac adverse effects of midostaurin^57^. Midostaurin has not been tested previously in cardiomyocyte hypertrophy to our knowledge. The mechanistic subnetwork analysis predicted that midostaurin inhibits TGFβ-induced hypertrophy via RAS and parallel MAP3K1/MAP3K2/3 pathways. Knockdown of these two nodes results in inhibition of TGFβ-induced hypertrophy in a manner comparable to the action of midostaurin (**Figure 4C**). Midostaurin is representative of two drugs in the model, midostaurin and loxapine, and these predictions would extend to other PKC inhibitors with the same binding characteristics.

### Experimental validation of context-specific drug predictions

Using our pharmacological model, we predicted that midostaurin would inhibit TGFβ-induced hypertrophy in a dose-dependent manner, but not norepinephrine-induced hypertrophy (**Figure 5A**). To experimentally validate these predictions, we tested the effects of midostaurin on cardiomyocyte hypertrophy in neonatal rat cardiomyocytes. Consistent with the model predictions, midostaurin inhibited TGFβ-induced hypertrophy in a dose-dependent manner, while midostaurin did not inhibit norepinephrine-induced hypertrophy (**Figure 5B**). The predicted inhibition of hypertrophy by midostaurin with TGFβ but not norepinephrine is also markedly apparent in the microscopy images (**Figure 5C**). Cell counts were based on DAPI staining of cardiomyocytes and reveal no significant difference between conditions (**Figure S5**). Taken together, these data support midostaurin as an antihypertrophic drug in cardiomyocytes.

**Figure 5.**
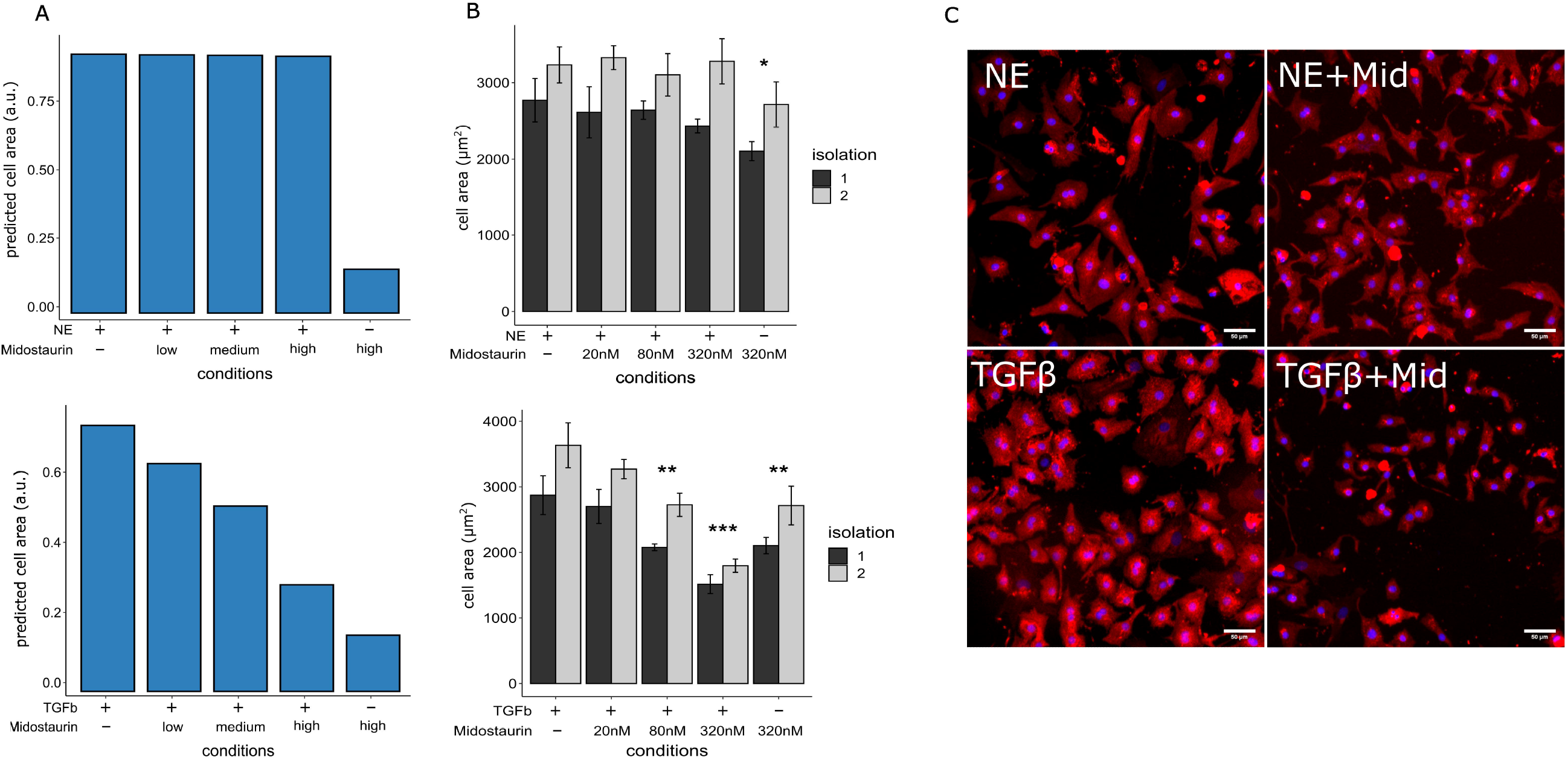
The model accurately predicts the differential effects of midostaurin on NE and TGFβ-induced cardiomyocyte hypertrophy. A) Predicted cell area response to midostaurin in TGFβ or NE-induced hypertrophy. B) Cell area response to varying doses of midostaurin treatment measured in two separate isolations of neonatal rat cardiomyocytes. C) Representative images of α-actinin stained cardiomyocytes stimulated by either TGFβ or NE and treated with 320 nm midostaurin. Two cell isolations with N = 4 wells each, error bar indicates SEM, * p<0.05, ** p<0.01, and *** p<0.001 comparing to stimulated condition without drug, 2-way ANOVA with Dunnet’s post-hoc correction.

### Modeling of polypharmacology and drug combinations

Polypharmacology and combination drug therapy provide opportunity for improved disease treatment^58–60^. Utilizing drugs with different targets improves the effect of the treatment. This may allow for a lower dose of each drug to be used, minimizing deleterious effects seen at higher doses. Likewise, using a drug that targets multiple pathways of disease may be more effective as a treatment. Using a polypharmacological approach will also reveal new off-target effects for existing drugs.

Zanubrutinib is representative of BTK inhibitors which have several off-target effects, including EGFR, ERBB and JAK inhibition^61^. Although these off-target effects introduce several toxicities in the context of cancer treatment, their role in cardiomyocyte hypertrophy is not well characterized^56^. Therefore, we simulated zanubrutinib activity in the context of neuregulin-induced hypertrophy, where zanubrutinib inhibits EGFR, ERBB and JAK simultaneously. The combined result of the inhibition of these nodes is an inhibition of hypertrophy (**Figure 6A**). We hypothesized that inhibition of multiple nodes may allow zanubrutinib to strongly inhibit hypertrophy regardless of the stimulus (**Figure 2A**).

**Figure 6.**
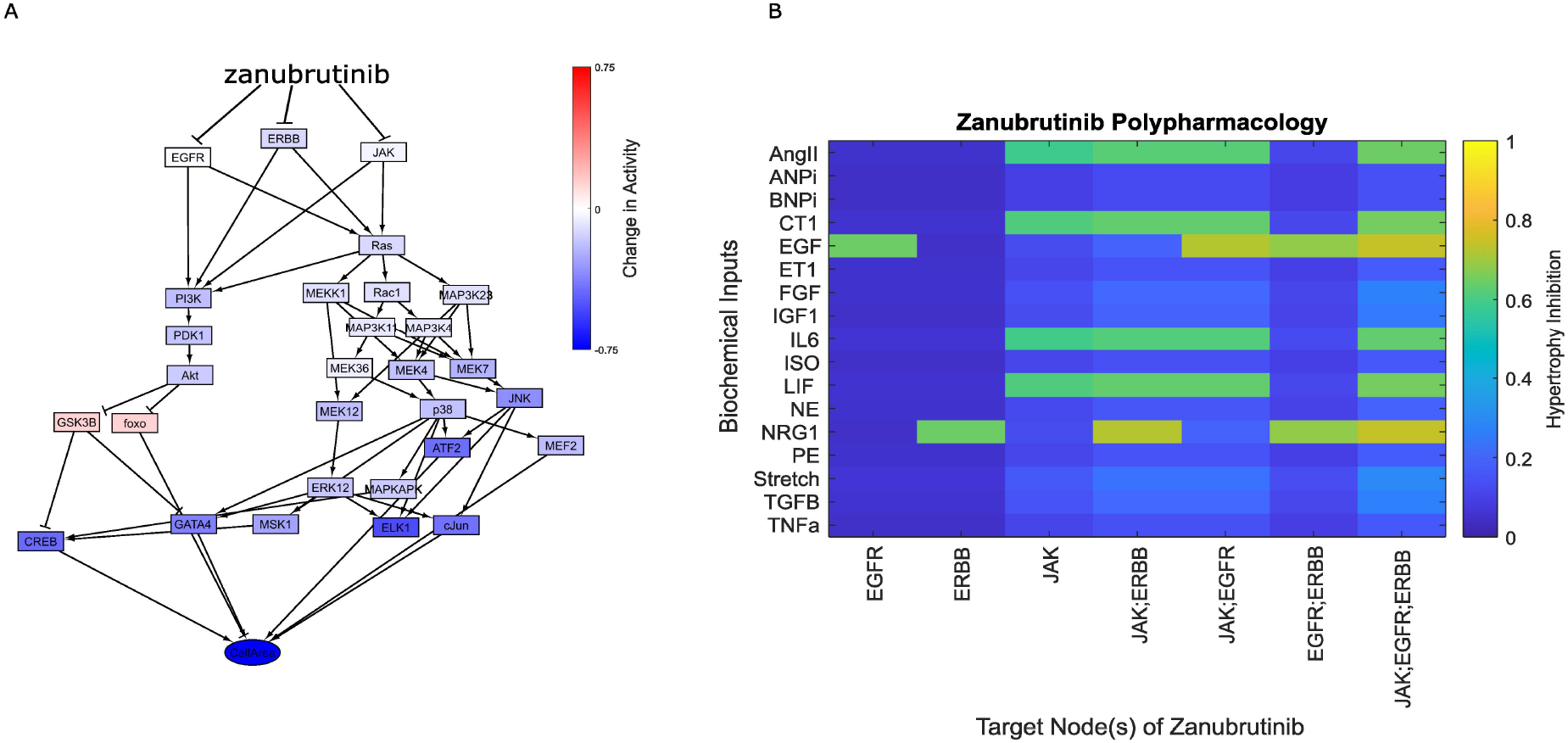
Zanubrutinib exhibits polypharmacological activity within the hypertrophy network. A) Zanubrutinib targets three distinct network nodes resulting in inhibition of neuregulin-induced hypertrophy. B) Simulation of the effects of zanubrutinib when accounting for the targets EGFR, ERBB, and JAK individually and combined.

Isolating the action of zanubrutinib on select targets reveal the inhibitory contribution of each node. When hypertrophy is induced with neuregulin, the model predicts that ERBB knockdown alone is sufficient to inhibit hypertrophy (**Figure 6B**). This inhibitory response is enhanced by the knockdown of JAK. EGFR knockdown inhibits EGF-induced hypertrophy, which is also enhanced by the knockdown of JAK. Knockdown of JAK alone is inhibitory of hypertrophy induced by LIF, IL6, CT1, and AngII. Zanubrutinib targeting of combinations of ERBB, EGFR, and JAK was predicted to be additive across conditions. Thus, the polypharmacology of zanubrutinib explains its robust capability to inhibit cardiomyocyte hypertrophy in a largely context-independent manner.

To investigate the effects of drug combinations on cardiomyocyte hypertrophy, pairs of drugs were simultaneously simulated. Combination responses of select representative drugs were compared to examine synergy in cell area, as quantified by Bliss independence. Drug combinations having a positive synergy score have cooperative inhibitory effects on hypertrophy while combinations with a negative synergy score have counteracting effects. A number of drugs were predicted to exhibit strong cooperative effects (**Figure S6**, **Figure 7A**). Brigatinib and irbesartan each inhibit hypertrophy alone, however combining the drugs results in an improved inhibitory effect (**Figure 7B**). Arsenic trioxide acts antagonistically towards brigatinib, and together results in a lack of hypertrophy inhibition.

**Figure 7.**
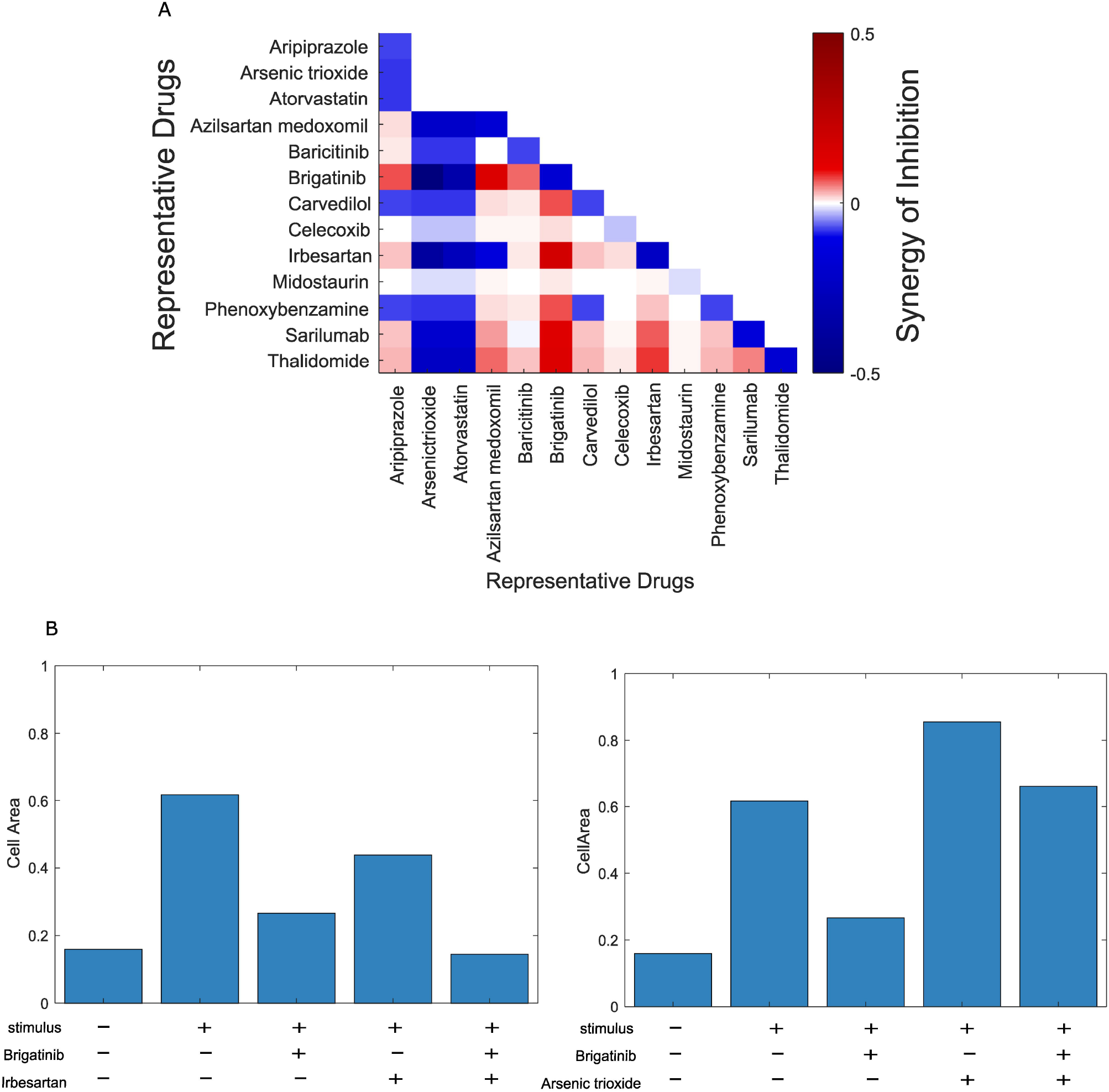
Selected drug pairs synergize in the inhibition of hypertrophy. A) Heatmap representing synergy scores in the inhibition of CM hypertrophy using the difference of percent inhibition and Bliss independence for drug pairs. Scores above 0 indicate drugs that act synergistically to inhibit hypertrophy. B) Brigatinib and irbesartan combine to improve the inhibition of hypertrophy. C) Brigatinib and arsenic trioxide act antagonistically resulting in a lack of hypertrophic inhibition.

## DISCUSSION

Cardiomyocyte hypertrophy develops as a result of distinct stimuli acting through several different signaling pathways^62^. Here, we have developed a pharmacological model of cardiomyocyte hypertrophy that predicts the effects of drugs validated with both *in vitro* and *in vivo* experiments. Our virtual drug screen identified 52 representative drugs that alter cardiomyocyte cell area. Our model predicts that different hypertrophic environments impact the effectiveness of drugs on inhibiting cardiomyocyte hypertrophy. This was supported by the prediction and experimental validation of midostaurin as an effective inhibitor of TGFβ-, but not NE-, induced hypertrophy. Using the network structure and virtual knockout screens, we identified subnetworks that describe antihypertrophic drug mechanisms. We investigated the polypharmacology of zanubritinib to explain how targeting three nodes broadens its efficacy across hypertrophic contexts. Finally, we used combinatorial drug analysis to identify synergism between brigatinib and irbesartan.

Celecoxib is a COX-2 inhibitor used to treat inflammation from rheumatoid arthritis and osteoarthritis^54^. Celecoxib has been shown to inhibit cardiomyocyte hypertrophy, and several mechanisms have been proposed to explain this^54,55,63^. 2,5-Dimethylcelecoxib, a celecoxib derivative that does not inhibit COX-2, has recently been shown to inhibit hypertrophy through the Akt pathway^32^. Our model supports that celecoxib inhibits isoproterenol-induced hypertrophy through a non-COX-2 mechanism. Additionally, we predicted that the efficacy of celecoxib is context dependent, with the greatest inhibitory effect occurring in the context of norepinephrine.

Midostaurin is a FLT3 inhibitor used to treat acute myeloid leukemia (AML). Various clinical trials emphasize the safety of midostaurin as a chemotherapeutic^64^. This drug was originally developed as a PKC inhibitor, and it potently inhibits αPKC in addition to FLT3 and KIT^65^. This is significant as αPKC has been shown to be involved in cardiomyocyte hypertrophy^66^. We show for the first time the effectiveness of midostaurin in hypertrophic inhibition, and further display the context-dependence of midostaurin. The successful inhibition of hypertrophy implicates midostaurin as a novel repurposed drug for reducing TGFβ-induced hypertrophy.

Combination therapeutics is challenging due to the complexity that arises from the sheer number of combinations. Signaling network models allow for the systematic analysis of these combinations^67^, improving therapeutic development for both novel and repurposed drugs. Using our network model, we predicted synergistic behavior between brigatinib and irbesartan in the inhibition of cardiomyocyte hypertrophy. Brigatinib, an inhibitor of both anaplastic lymphoma kinase inhibitor (ALK) and epidermal growth factor receptor (EGFR), has been shown to be effective in ALK-positive non-small-cell lung cancer^68,69^. Irbesartan is an angiotensin receptor blocker (ARB) used for hypertension and known to inhibit hypertrophy^70^. Brigatinib shows minimal cardiotoxicity compared to other ALK-tyrosine kinase inhibitors^71^, but has not been tested in the context of hypertrophy. Although each drug alone is predicted to inhibit hypertrophy to some extent, the combination of both drugs predicts an improved outcome. Further experimentation will be needed to validate these predictions.

There are several limitations to the current study, the primary one being the incompleteness of the hypertrophy model due to limited data availability. The signaling networks of cardiomyocyte hypertrophy are well characterized^5^, but not all-inclusive. Several drug predictions that contradict prior literature results focus on the role of HDAC signaling in hypertrophy. The model does not currently explore the complexity of HDAC activity in cardiomyocyte hypertrophy, and further work would require investigation of this pathway. Drug behavior was modeled using binding characteristics but did not consider target affinities. While the current network model accurately predicts *in vivo* cardiac hypertrophy for 52 transgenic mice experiments^72^, performing drug studies *in vivo* would further validate the predicted behavior of antihypertrophic drugs. The hypertrophic stimuli for this study were simulated individually, whereas clinical states of hypertrophy would best be represented by a combination of stimuli. Future work using this model could use these input combinations to mimic *in vivo* and clinical scenarios. This model is only able to predict mechanisms of drugs that target nodes included in the network. Further work can be done to expand the network to allow for investigation of drugs that inhibit hypertrophy through alternate regulators.

The development of computational methods to screen for antihypertrophic drugs is critical in the prevention of heart failure. Towards this goal, we developed a pharmacological model of cardiomyocyte hypertrophy. This model provides a new tool for identifying antihypertrophic drugs and their network mechanisms.

## Acknowledgments

We thank Anirudha Chandrabhatla, Anders Nelson, Tim McKinsey, Ali Khalilimeybodi, Bryana Harris, and Mukti Chowkwale for their contributions to this work.

## Data and materials availability

All data are available in the main text or the supplementary materials.

