## Supplementary Materials for "Virtual drug screen reveals context-dependent inhibition of cardiomyocyte hypertrophy"

**Title:**

**This PDF file includes:**

Table S1

Figs. S1 to S6

**Supplementary Materials:**

**Table S1.** Representative drugs identified from the virtual screen along with their characteristics

| **Drug** | **Binding** | **Action** | **Target** |
| --- | --- | --- | --- |
| 4-Methoxyamphetamine | Competitive | Agonist | aAR |
| AMG-510 | Non-Competitive | Antagonist | Ras |
| Acepromazine | Competitive | Antagonist | aAR |
| Acetylsalicylic acid | Non-Competitive | Antagonist | IkB;IKK;ET1R |
| Adapalene | Competitive | Antagonist | cJun |
| Afatinib | Non-Competitive | Antagonist | EGFR;ERBB |
| Alpelisib | Non-Competitive | Antagonist | PI3K |
| Alprenolol | Competitive | Antagonist | BAR |
| Ambrisentan | Competitive | Antagonist | ET1R |
| Angiotensin II | Competitive | Agonist | AT1R |
| Aprindine | Competitive | Antagonist | CaM |
| Arbutamine | Competitive | Agonist | BAR |
| Aripiprazole | Competitive | Antagonist | aAR;BAR |
| Arsenic trioxide | Competitive | Agonist | ERK12;HDAC;Akt;IKK;cJun |
| Atorvastatin | Competitive | Antagonist | HDAC |
| Azilsartan medoxomil | Competitive | Antagonist | AT1R |
| Baricitinib | Non-Competitive | Antagonist | JAK |
| Bosutinib | Non-Competitive | Antagonist | MAP3K23;CaMK;MEK12 |
| Brigatinib | Competitive | Antagonist | EGFR;ERBB;IGF1R |
| Calcium Phosphate | Competitive | Agonist | CaM |
| Carvedilol | Competitive | Antagonist | aAR;BNP;BAR |
| Celecoxib | Non-Competitive | Antagonist | PDK1 |
| Chlorpromazine | Competitive | Antagonist | aAR;CaM |
| Clenbuterol | Competitive | Agonist | TNFa;BAR |
| Cobimetinib | Non-Competitive | Antagonist | MEK12 |
| Dabrafenib | Non-Competitive | Antagonist | Raf1 |
| Dacomitinib | Non-Competitive | Antagonist | EGFR |
| Dipivefrin | Competitive | Agonist | aAR;BAR |
| Ellagic acid | Non-Competitive | Antagonist | PKA;PKC |
| Epinephrine | Competitive | Agonist | aAR;TNFa;BAR |
| Erdafitinib | Non-Competitive | Antagonist | FGFR |
| Foreskin keratinocyte (neonatal) | Competitive | Agonist | EGFR |
| Fostamatinib | Non-Competitive | Antagonist | MEK5;PDK1;mTor;IKK;FAK;ERK5;PKA;PKD;  EGFR;MEK36;MAP3K23;MAP3K11;MAPKAPK;  CaMK;p38;MEKK1;GSK3B;MAP3K4;ERBB;JAK;  Raf1;FGFR;PI3K;MEK12;PKC |
| Irbesartan | Competitive | Antagonist | cJun;AT1R |
| Isoprenaline | Competitive | Agonist | BAR;ERK12 |
| Lapatinib | Non-Competitive | Antagonist | ERBB;EGFR |
| Linsitinib | Non-Competitive | Antagonist | IGF1R |
| Lithium cation | Non-Competitive | Antagonist | GSK3B |
| Mecasermin | Competitive | Agonist | IGF1R |
| Midostaurin | Non-Competitive | Antagonist | PKC |
| Pazopanib | Non-Competitive | Antagonist | FGF;FGFR |
| Pertuzumab | Non-Competitive | Antagonist | ERBB |
| Phenoxybenzamine | Competitive | Antagonist | aAR;CaM;BAR |
| Pseudoephedrine | Competitive | Agonist | aAR;NFkB;TNFa;BAR;ATF2 |
| Purvalanol | Non-Competitive | Antagonist | ERK12 |
| Regorafenib | Non-Competitive | Antagonist | Raf1;p38;FGFR |
| Resveratrol | Non-Competitive | Antagonist | Akt |
| Sarilumab | Competitive | Antagonist | IL6R |
| Sorafenib | Non-Competitive | Antagonist | Raf1;FGFR |
| Terazosin | Competitive | Antagonist | aAR;TGFB |
| Thalidomide | Competitive | Antagonist | NFkB;FGFR;TNFa |
| Zanubrutinib | Non-Competitive | Antagonist | JAK;EGFR;ERBB |
| Vinorelbine | Non-Competitive | Antagonist | p38 |

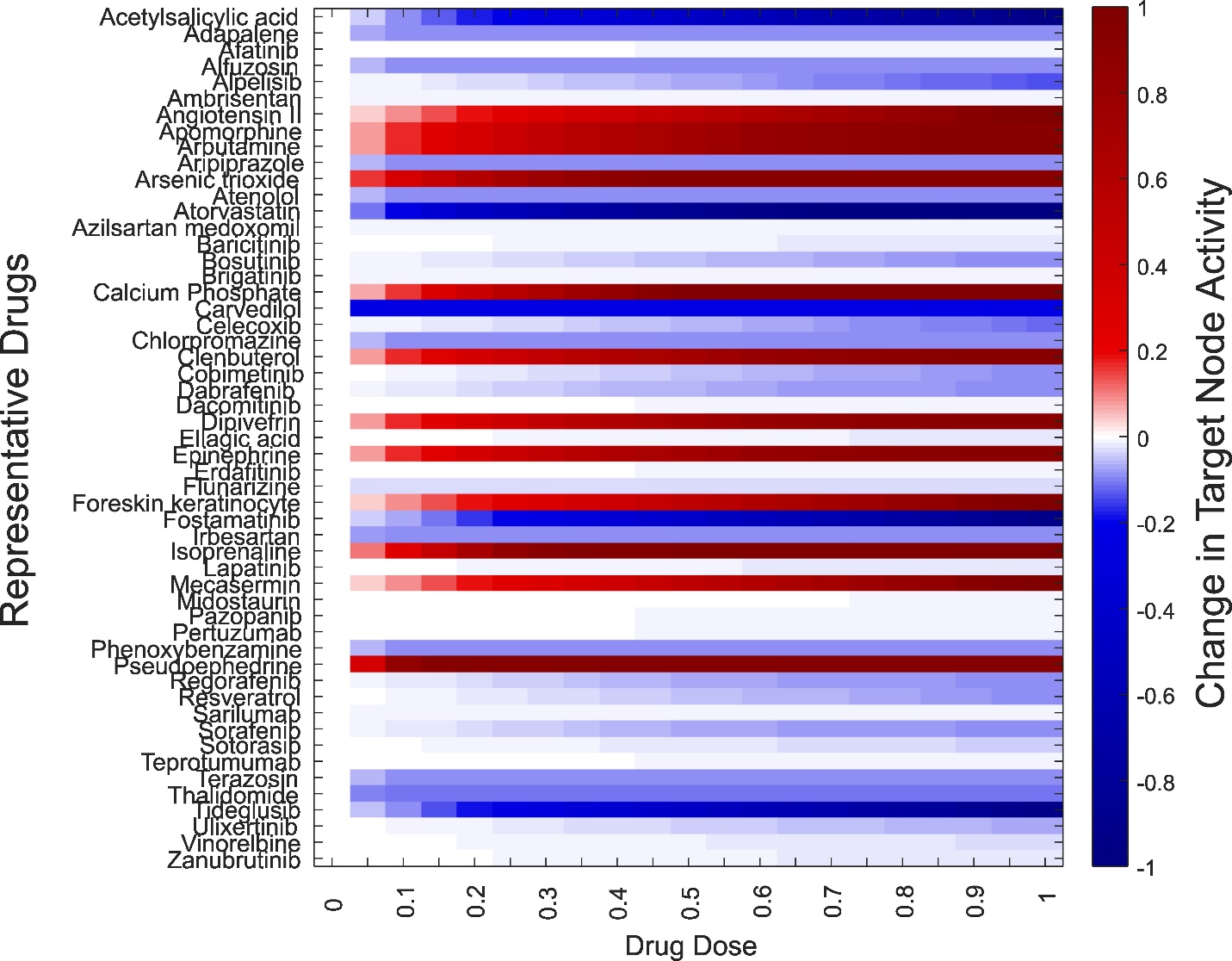
**Figure S1. Simulation of the drug target dose response.** Heatmap represents the response to a drug acting on its target node. If the drug has multiple targets, only the first node’s response is shown.

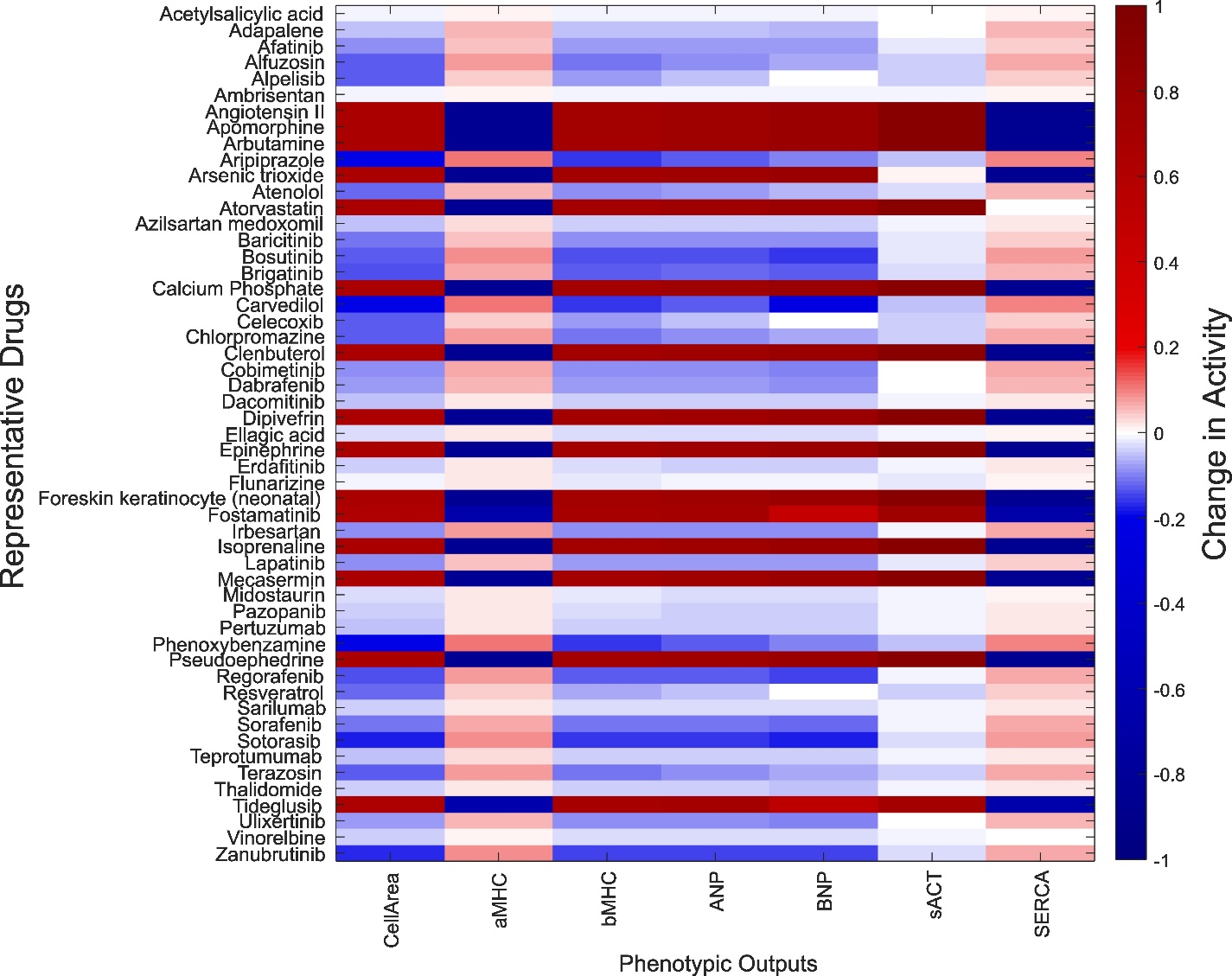
**Figure S2. Simulation of the drug effect on phenotypic outputs from angiotensin II-induced hypertrophy.** The predicted effect of each drug on the 7 phenotypic outputs is represented as the change in activity of that output.

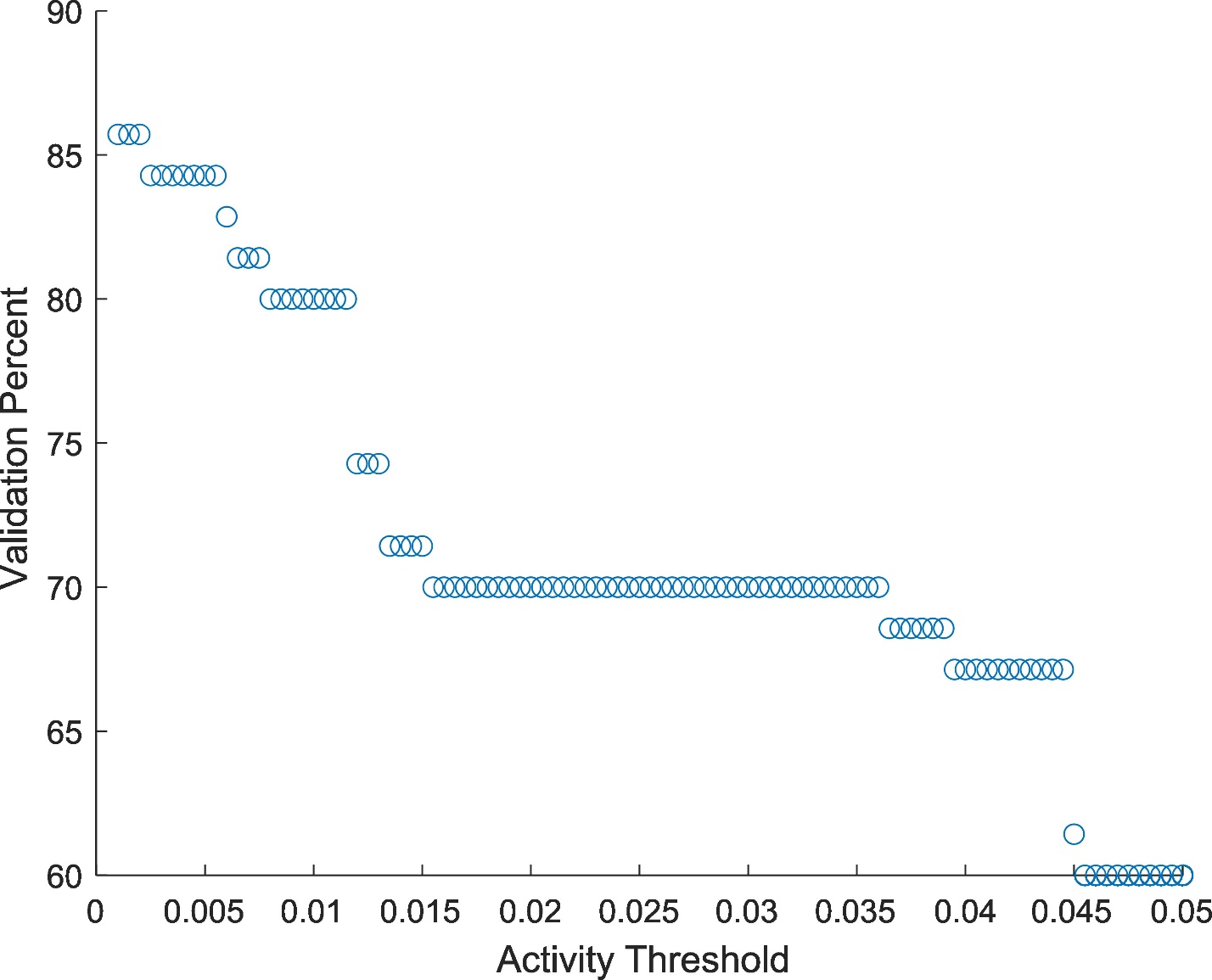
**Figure S3. Model validation is robust across large range of threshold values.** Validation percent of the cardiomyocyte hypertrophy model is shown for activity threshold levels between 0.001 and 0.05.

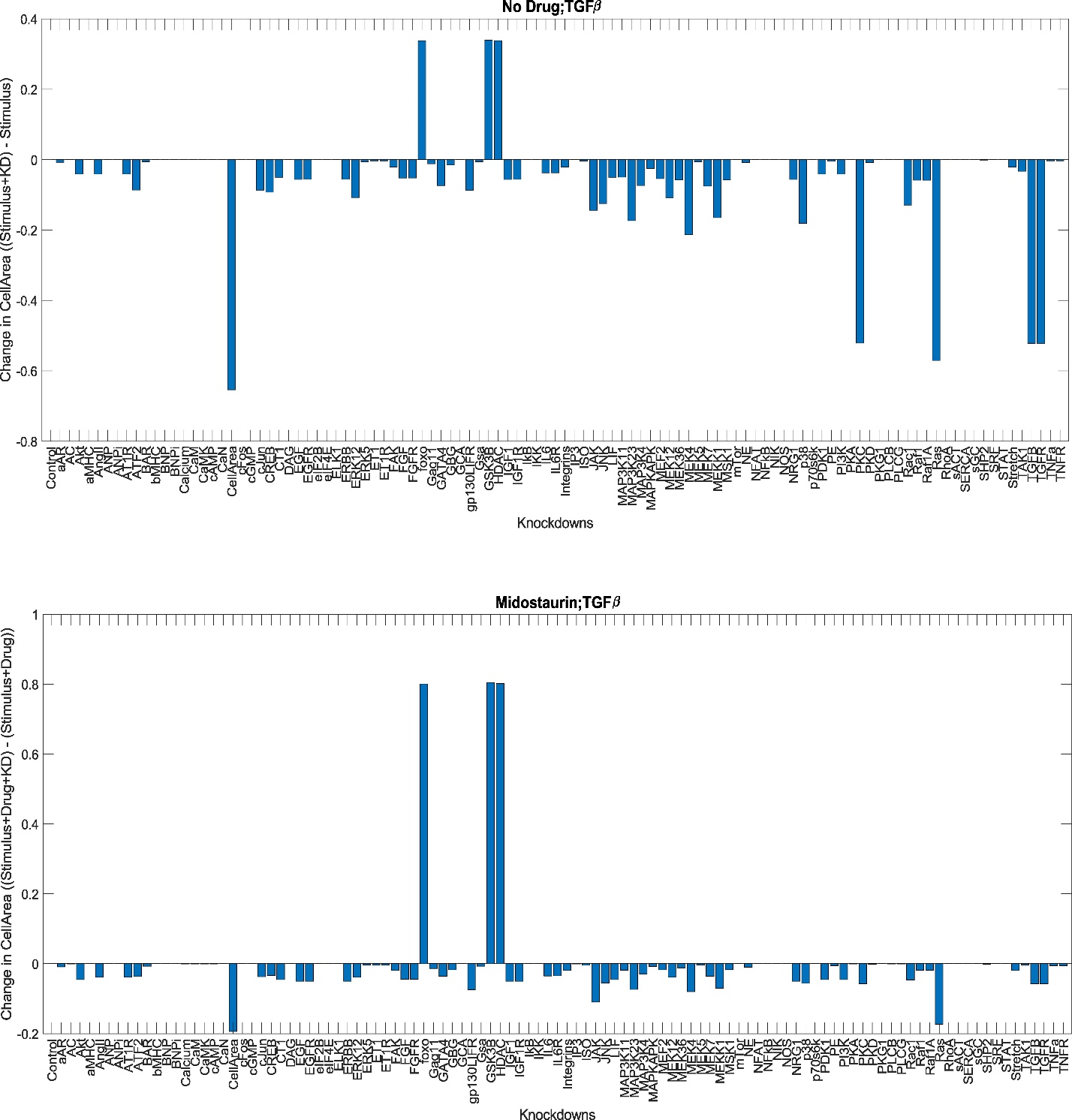
**Figure S4. Knockdown screen identifies nodes that facilitate drug activity.** A) The difference in TGFβ-stimulated cell area with and without individual node knockdowns. B) The difference in midostaurin treated, TGFβ -stimulated cell area with and without individual node knockdowns. Taking the difference of A and B results in the identification of nodes that are essential for the action of midostaurin in TGFβ -induced hypertrophy.

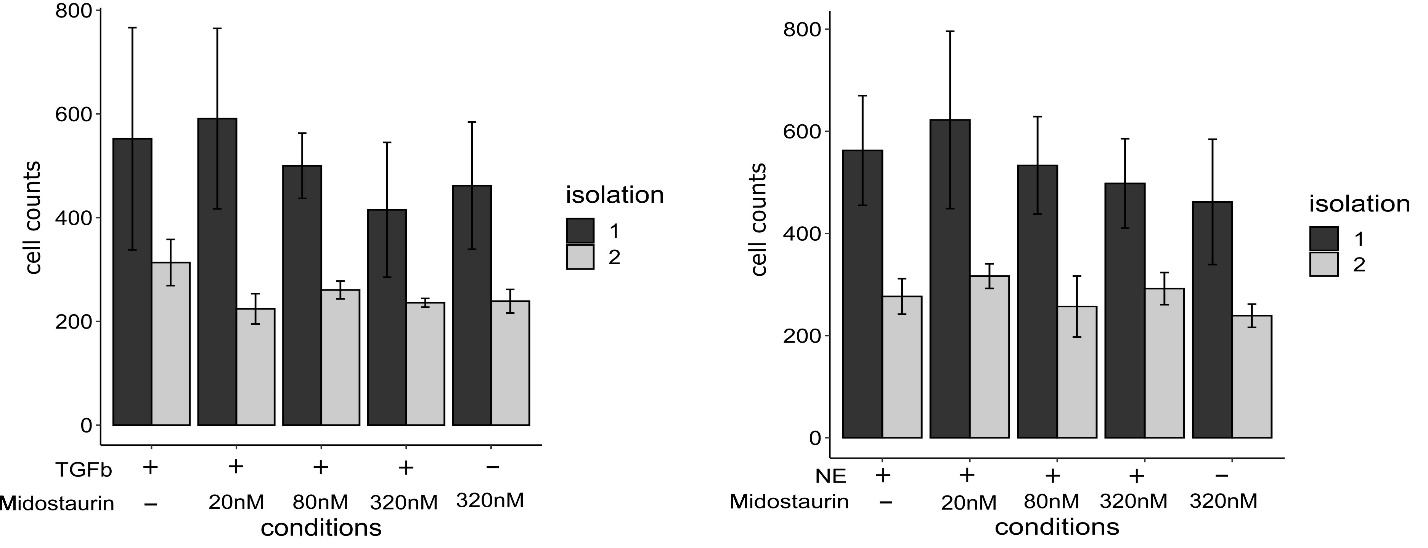
**Figure S5. Cardiomyocyte viability is not significantly decreased by midostaurin.** Cell counts were measured based on DAPI staining for two separate cardiomyocyte isolations. No significant difference was seen compared to the presence of hypertrophic stimulus alone, based on 2-way ANOVA with Dunnet’s post-hoc correction.

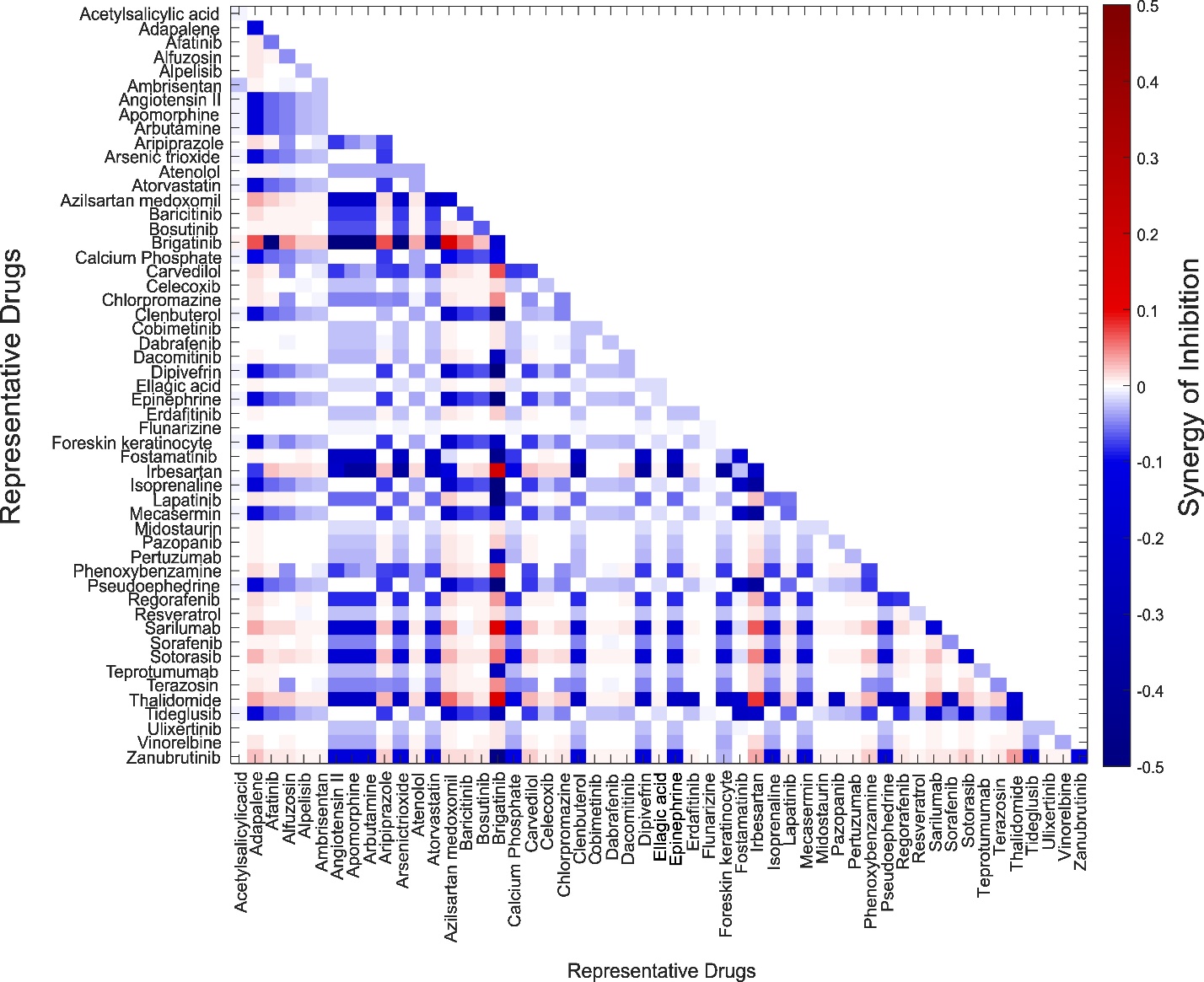
**S6. Synergy scores for pairwise combinations of representative drugs in the inhibition of cardiomyocyte hypertrophy.** Heatmap representing synergy scores in the inhibition of CM hypertrophy using Bliss independence for drug pairs. Scores above 0 indicate drugs that act synergistically to inhibit hypertrophy.
